# Efferent control of hearing sensitivity and protection via inner ear supporting cells

**DOI:** 10.1101/2020.12.03.409946

**Authors:** Hong-Bo Zhao, Li-Man Liu, Ning Yu, Yan Zhu, Ling Mei, Jin Chen, Chun Liang

## Abstract

It is well-known that medial olivocochlear (MOC) efferent nerves innervate outer hair cells (OHCs) and inhibit OHC electromotility to control hearing sensitivity. Here, we report that the MOC nerve fibers associated with MOC neurotransmitter acetylcholine (ACh) receptors and presynaptic vesicular acetylcholine transporters (VAChT) also innervated the cochlear supporting cells (SCs). Application of ACh could evoke inward currents in SCs and reduced gap junctions (GJs) between them with long-lasting effect. We further found that uncoupling of GJs between SCs shifted OHC electromotility to the left hyperpolarization direction and enhanced the direct effect of ACh on OHC electromotility. Diminution of this control by deficiency of GJs between SCs compromised the regulation of active cochlear amplification and increased susceptibility to noise. These data suggest that the MOC efferent system has functional innervations with SCs. Such efferent innervation may have an important role in the protection of hearing from noise and slow efferent effect.

## Introduction

The cochlea is the auditory sensory organ in mammals. In addition to ascending afferent auditory nerves (ANs) projecting to the brain, the cochlea also receives descending efferent nerve fibers to hair cells forming a negative feedback loop to control hair cell activity (Guinan, 2006). The cochlear efferent nerves are composed of the medial olivocochlear (MOC) nerve fibers, which project to outer hair cells (OHCs), and the lateral olivocochlear (LOC) nerve fibers, which project to inner hair cells (IHCs) and form synapses with the dendrites of type-I afferent ANs under IHCs (Maison et al., 2003; Guinan, 2006; Lustig, 2006). MOC nerve fibers are cholinergic and acetylcholine (ACh) is a primary neurotransmitter, while LOC nerves have cholinergic and dopaminergic fibers, releasing ACh, dopamine, and other neurotransmitters (Eybalin, 1993; Lustig, 2006; Maison et al., 2012). The cochlear efferent system plays an important role in many aspects of hearing, such as the regulation of hearing sensitivity, the discrimination of sounds in background noise, and the protection of hearing from acoustic trauma (Guinan, 2006; Lustig, 2006; Clause et al., 2017; Boero et al., 2018).

The cochlea contains sensory hair cells and non-sensory supporting cells (SCs). SCs in the cochlea provide supporting function to hair cell activity. For example, Deiters cells (DCs) and pillar cells (PCs) in vicinity of OHCs act as a scaffold to support OHCs standing on the basilar membrane allowing OHC motility amplifying sound stimulation induced the basilar membrane vibration to increase hearing sensitivity and frequency selectivity (Brownell et al., 1985; Zhao and Santos-Sacchi, 1999; Ashmore, 2008). This prestin-based active cochlear amplification or mechanics is required for mammalian hearing. Deficiency of this active cochlear amplification can induce hearing loss (Zheng et al., 2000; Liberman et al., 2002). In anatomy, cochlear SCs are extensively coupled by gap junctions (GJs) (Kikuchi et al., 1995; Forge et al., 2003; Zhao and Yu, 2006). Connexin 26 (Cx26) and Cx30 are predominant GJ isoforms in the cochlea (Forge et al., 2003; Zhao and Yu, 2006; Liu and Zhao, 2008). However, there is neither connexin expression in hair cells nor GJs between hair cells and SCs (Kikuchi et al., 1995; Zhao and Santos-Sacchi, 1999; Zhao and Yu, 2006; Yu and Zhao, 2009). Cx26 mutations can cause hearing loss, responsible for >50% of nonsyndromic hearing loss (Castillo and Castillo, 2011; Chan and Chang, 2014; Wingard and Zhao, 2015). Our previous studies also demonstrated that SCs and SC GJs can modulate OHC electromotility and participate in active cochlear amplification (Yu and Zhao, 2009; Zhu et al., 2013, Zong et al., 2017). These data suggest that SCs in the cochlea have important roles in hearing. However, the detailed SC function still remains largely undetermined.

For example, it was reported that the cochlear SCs have nerve innervations (Wright and Preston, 1976; Stopp and Comis, 1979; Liberman et al., 1990; Nadol and Burgess, 1994; Burgess et al., 1997; Bruce et al., 2000). However, the source and function of these nerve innervations in the SCs remain unclear or are under debated (Fechner et al., 1998, 2001). In this study, we found that MOC efferent nerves had innervation with SCs to regulate GJs between them, which could enhance the direct effect of ACh on OHC electromotility. Diminution of this SC-mediated efferent control compromised the regulation of active cochlear amplification and the protection of hearing from noise trauma.

## Results

### MOC innervation in the cochlear SCs

Efferent MOC nerves are well-known to project to OHCs in the cochlea. However, we found that the MOC fibers also have branches projecting to the cochlear SCs. Fig. 1 shows that the MOC fibers, which were labeled by Tuj1 or neurofilament (NF), passed through the cochlear tunnel projecting to SCs in addition to OHCs. The MOC nerves had branches projecting to Deiters cells (DCs), outer pillar cells (OPCs), and Hensen cells (HCs) (Fig. 1A-F). The nerve branch projected from the first row of OHCs to the second and third row of DCs (indicated by white arrowheads in Fig. 1C). The MOC nerves are cholinergic fibers and ACh is a primary neurotransmitter. Besides intensive labeling for ACh receptors (AChRs) at the basal pole of OHCs, the positive AChR labeling was visible at the SCs with the MOC nerves (indicated by white arrowheads in Figs. 1D&E and 1-S1) but not with type-II AN fibers (Fig. 1-S1A&B). To further immunostain the synaptic terminals, we also used antibody to presynaptic vesicular acetylcholine transporter (VAChT) to show presynaptic endings (Maison et al., 2003; Webber et al., 2021). Immunofluorescent staining for VAChT shows that besides large puncta of labeling at the OHC basal pole, small puncta of labeling associated with MOC nerves were visible at SCs (indicated by white arrowheads in Fig. 1A&B). Such VAChT labeling also was not associated with eGFP-targeted type II AN fibers (Fig. 1A&B). In addition, MOC nerves and type II AN fibers showed different locations to cross the cochlear tunnel. Tuj1-labeled MOC fibers crossed the cochlear tunnel at the middle level of the tunnel projecting to OHCs and SCs, whereas eGFP-targeted type II AN fibers passed through the cochlear tunnel along its bottom then rose up projecting to OHCs (Fig. 1A&B).

**Fig. 1.**
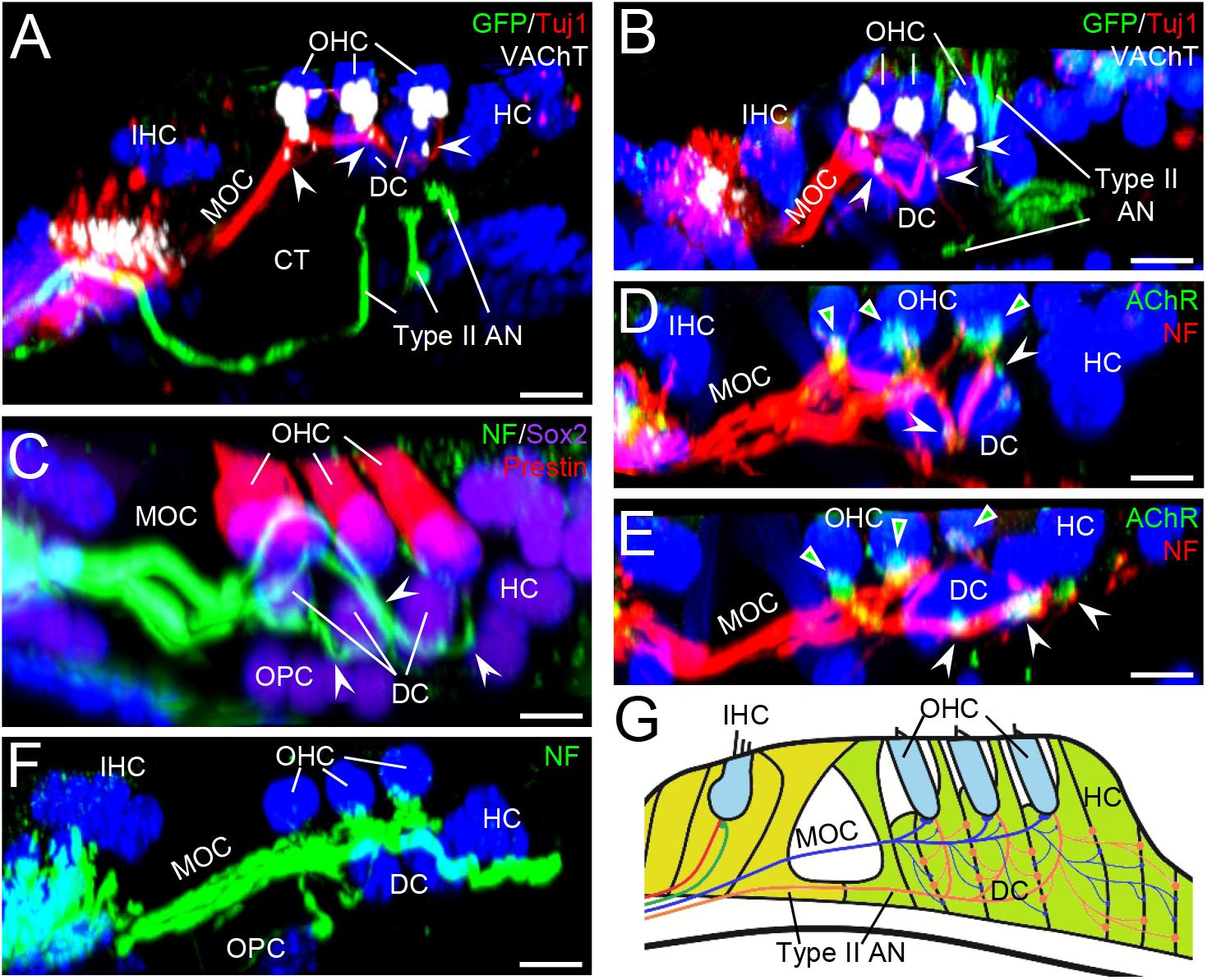
Innervation of the medial olivocochlear (MOC) efferent nerves in the cochlear OHCs and supporting cells (SCs) in mice. **A-B:** Immunofluorescent staining of the cochlear sensory epithelium of peripherin-eGFP transgenic mice (P33 old) for eGFP (green), Tuj1 (red), and vesicular acetylcholine transporter (VAChT, white). Type II auditory nerves (ANs) in peripherin-eGFP transgenic mice were labeled by eGFP, which was further enhanced by immunofluorescent staining for eGFP. The images were the cross-section view of 3D images constructed from z-stack of confocal scanning images. White arrowheads indicate VAChT labeling at Deiters cells (DCs) with MOC fibers. IHC: inner hair cell, OHC: outer hair cell, HC: Hensen cell, and CT: the cochlear tunnel. **C:** Projections of MOC fibers to DCs, HCs, and outer pillar cells (OPCs). The cochlear sensory epithelium was stained for neurofilament (NF, green), prestin (red), and Sox2 (purple) for labeling nerve fibers, OHCs, and SCs, respectively. White arrowheads indicate the MOC branch projecting to the DC, HC, and PC, respectively. **D-E:** ACh receptor (AChR) expression at the OHCs and SCs. AChR and NF are labeled by green and red colors, respectively. Triangles with green color indicate AChR expression at the OHC basal pole and white arrowheads indicate expression of AChR at DCs and HCs. **F:** Projections of MOC fibers to DCs, HCs, and OPCs. **G:** Schematic drawing of innervations of MOC and type II AN fibers in OHC and SC areas in the cochlea. Scale bars: 10 µm.

### Responses to ACh in OHCs and SCs

To further test functional innervation, we recorded responses of OHCs and SCs to MOC neurotransmitter ACh. ACh could evoke a typical inward current in OHCs and SCs (Fig. 2B&D). At holding at -80 mV, 0.1 mM ACh could evoke -0.31±0.07 nA (n=12) and -0.25±0.07 nA (n=5) inward currents in OHCs and DCs, respectively, in guinea pigs (Fig. 2F). There was no significant difference between them (P=0.60, two-tail t test). As ACh was repeatedly applied, the evoked inward current could be repeatedly recordable (Fig. 2B&D). We measured ACh dose-curve in single DCs in the guinea pig (Fig. 2E). The EC_50_ was 72.6 μM and Hill’s coefficient was 1.64 (Fig. 2E). The ACh-evoked inward current was also visible in mouse SCs (Fig. 2-S1). However, comparing with the responses in guinea pigs, the ACh-evoked current in mouse SCs appeared small. At -80 mV, 0.5 mM ACh evoked currents in guinea pig and mouse DCs were -0.38±0.07 nA (n=12) and -0.14±0.03 nA (n=7), respectively (Fig. 2-S1B). The evoked current in mouse DCs was significantly smaller than that in guinea pigs (P=0.006, two-tail t test).

**Fig. 2.**
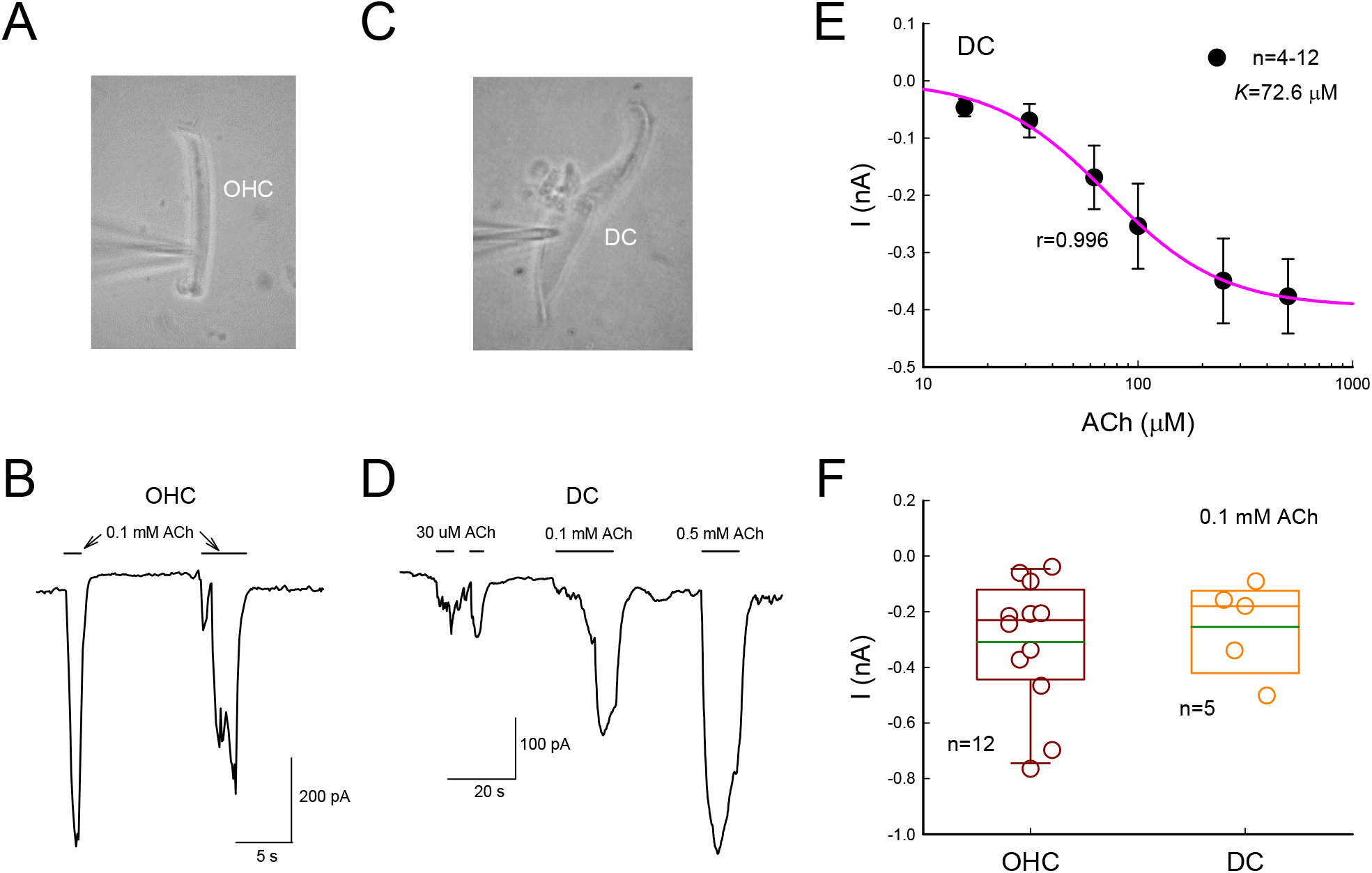
ACh-evoked inward current in OHCs and the cochlear SCs in guinea pigs. **A-B:** ACh-evoked inward current in OHCs. Panel A is a captured image of patch-clamp recording in single OHC. A horizontal bar in panel B represents application of 0.1 mM ACh. The membrane potential was clamped at -80 mV. **C-D:** ACh-evoked inward current in a DC. Horizontal bars represent the application of ACh. The membrane potential was clamped at -80 mV. **E:** Dose curve of ACh-evoked current in single DCs. The smooth line represents data fitting to a Hill’s function: *I=a* * *C^n^/(K^n^ + C^n^)*, where n=1.64 (Hill coefficient) and *K*=72.6 µM (EC_50_) for ACh. **F:** Inward currents measured in OHCs and DCs at -80 mV evoked by 0.1 mM ACh. Green lines in boxes represent mean levels, which are -0.31±0.07 and -0.25±0.07 nA in OHCs and DCs, respectively.

In the experiment, we found almost all (91-100%, i.e., 20/22 and 11/11) of recorded DCs and most (75-83%, i.e., 6/8 and 10/12) of the recorded HCs in guinea pigs and mice had responses to ACh (Fig. 2-S2). However, only few (1/6 and 0/4 in guinea pigs and mice, respectively) of recorded Claudius cells (CCs) had response to ACh (Fig. 2-S2). This corresponded well with observation in immunofluorescent staining that MOC fibers mainly innervated with DCs and HCs, and rarely with CCs in the cochlea (Fig. 1).

### Amplification of ACh response in SCs by GJs

SCs in the cochlea are extensively coupled by GJs, which provide an intracellular electrical conduit between cells to synchronize or amplify electrical responses in a cell group. Fig. 3 shows the ACh-evoked currents in 2DCs. At -80 mV, the current evoked by 0.5 mM ACh in 2DCs in guinea pigs was -1.44±0.76 nA (n=6), more than 3 times larger than that (-0.38±0.07 nA, n=12) in single DCs. In mice, the current evoked by 0.5 mM ACh in 2DCs was -0.29±0.14 nA (n=4), 2 times larger than that (0.14±0.03 nA, n=7) in in the single DCs.

**Fig. 3.**
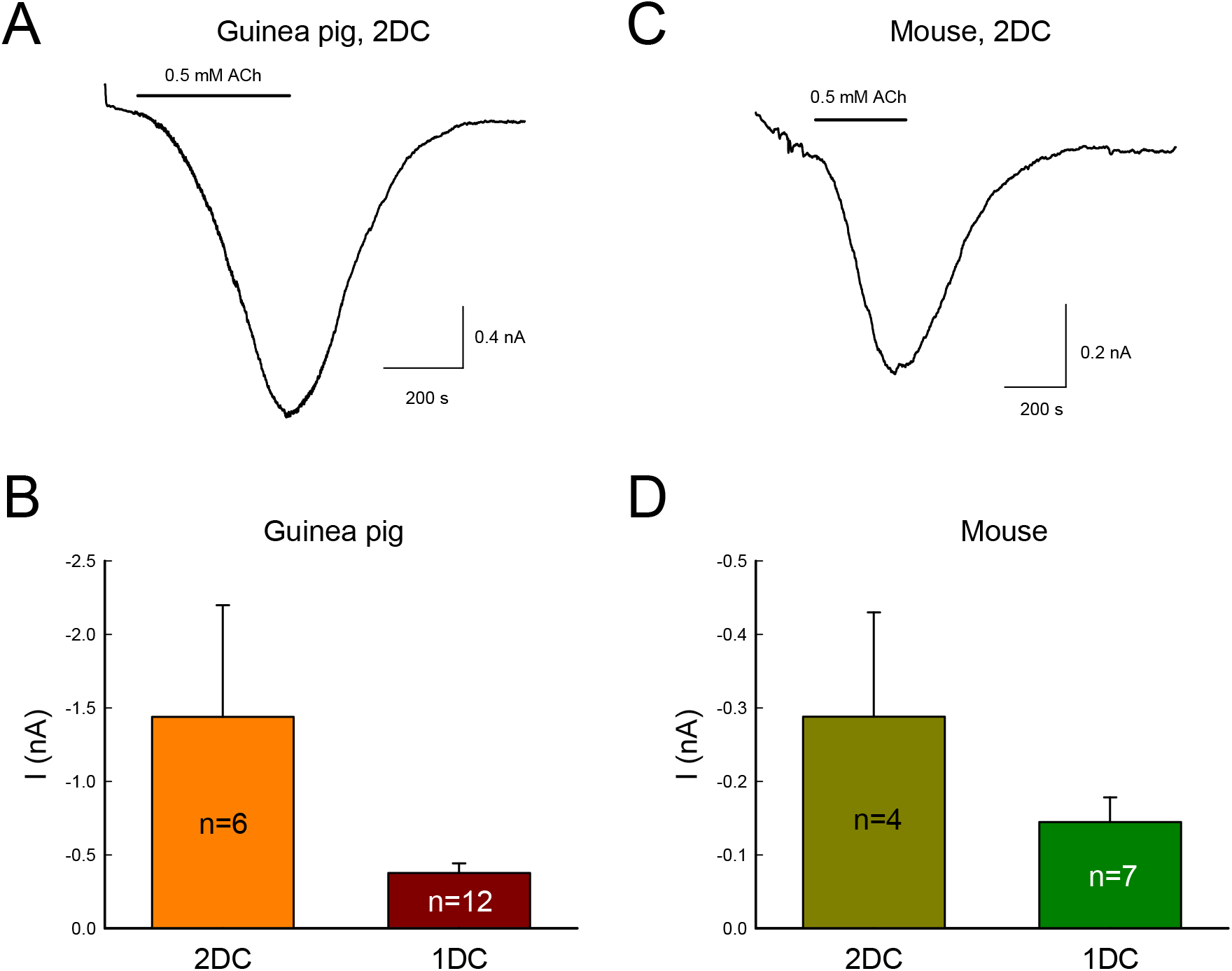
Amplification of ACh-evoked inward currents in SCs by gap junctional coupling. **A-B:** ACh-evoked current in 2DCs in guinea pigs. In comparison with 1DC, the current evoked by 0.5 mM ACh in 2DCs is -1.44±0.76 nA and more than 3 times larger than that (-0.38±0.07 nA) in single guinea pig DCs. **C-D:** ACh-evoked current in mouse DCs. In comparison with ACh-evoked current (-0.14±0.03 nA) in single DC in mice, the average current evoked by 0.5 mM ACh in mouse 2DCs is -0.29±0.14 nA and increased by 2 times.

### Effect of ACh on GJs between SCs

ACh receptors are permeable to Ca^2+^ ions to elevate intracellular Ca^2+^, which can close GJ channels, i.e., uncouple GJs (Spray et al., 1982; Sato and Santos-Sacchi, 1994). We further assessed the effect of ACh on GJs between SCs. As reported previously (Zhao and Santos-Sacchi, 1998; Zhu and Zhao, 2012), input capacitance (*C_in_*) was recorded to assess GJ coupling between the cochlear SCs. *C_in_* in the 2DCs and single DCs in guinea pigs was 69.9±3.62 pF (n=14) and 32.6±1.54 pF (n=24), respectively (Fig. 4-S1A). *C_in_* of single DCs was almost half of *C_in_* in 2DCs. Application of 0.5 mM ACh reduced *C_in_* in pairs of DCs to a half value (Fig. 4A&C), indicating that two coupled DCs were uncoupled, i.e., GJ channels became closed. In both guinea pigs and mice, *C_in_* of 2DCs was reduced to 69.3±7.94 % (n=8) and 64.0±10.7 % (n=6), respectively, under application of 0.5 mM ACh (Fig. 4-S1B). The uncoupling effect of ACh was reversible. After stop of perfusion of ACh, *C_in_* was increased and returned to the coupled levels (Fig. 4A&C). However, long-term application of ACh on the order of minutes usually caused the uncoupling irreversible (Fig. 4B&D). Fig. 4D showed uncoupling effect of ACh in a HC group. *C_in_* appeared a step-reduction during the long-term (∼ 8 min) treatment of 0.5 mM ACh. However, *C_in_* in single cell had no apparent change for application of ACh (Fig. 4E&F), although ACh also could evoke current response in the recording cell (data not shown). This further indicated that *C_in_* measurement was not dependent on changes in membrane current.

**Fig. 4.**
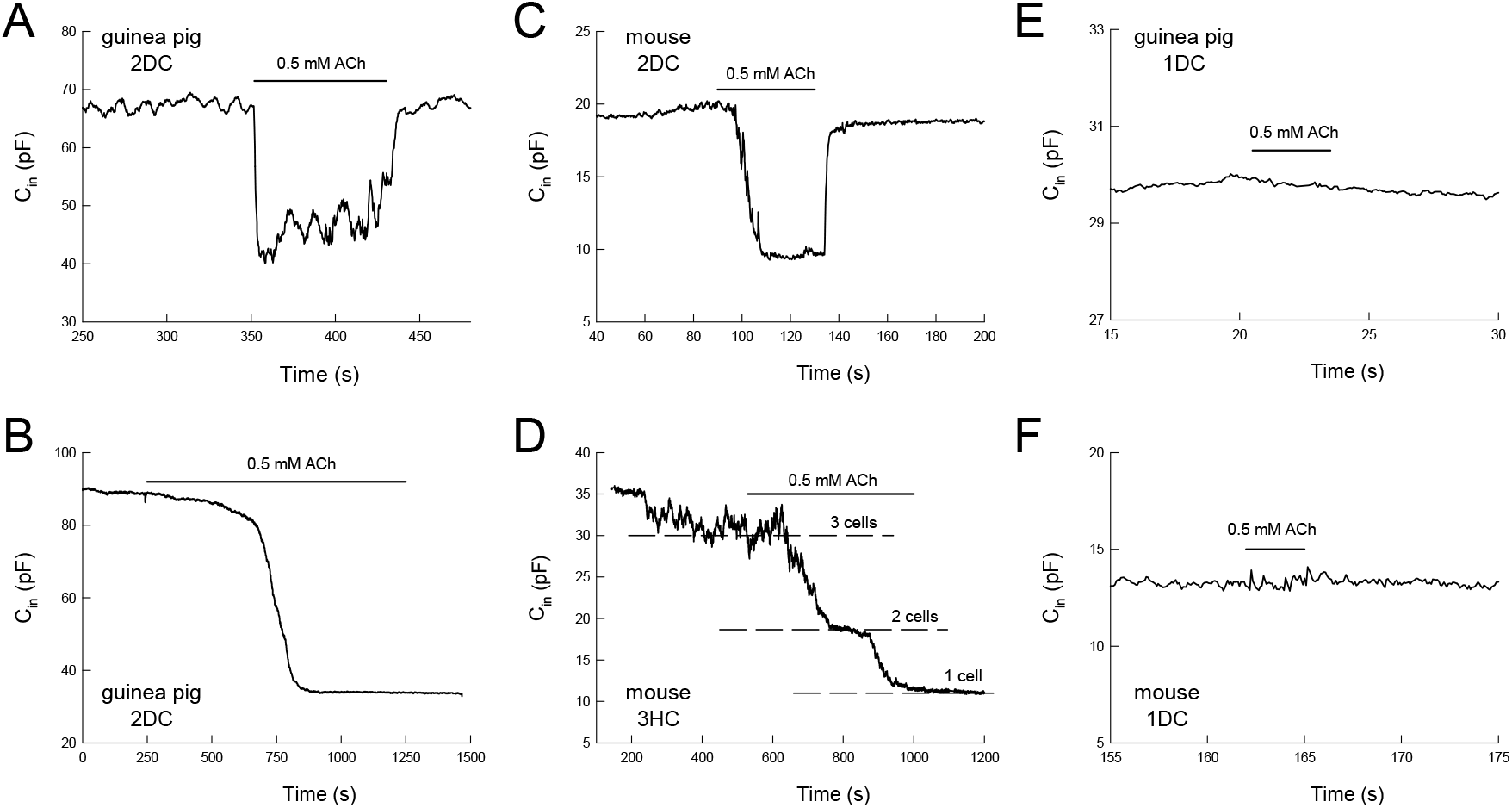
ACh-induced uncoupling effect on GJs between the cochlear SCs. The GJ coupling between the SCs was measured by input capacitance (*C_in_*). **A-B:** ACh-induced uncoupling effect on GJs between SCs in guinea pigs. Panel A shows the reversible uncoupling of GPs between 2DCs by ACh. After closing GJs between cells under treatment of 0.5 mM ACh, C_in_ was reduced to the half value to one cell level. Panel B shows uncoupling of GJs in long-term application of ACh. **C-D:** ACh-induced uncoupling effect on GJs between SCs in mice. Panel D shows step-changes in C_in_ for uncoupling of GJs among three-coupled HCs in long-term application of ACh. **E-F:** No apparent changes in C_in_ in single DC for the same ACh (0.5 mM) application in both guinea pigs and mice.

To further assess the effect of ACh on GJ function, we also used fluorescence recovery after photobleaching (FRAP) to measure GJ permeability (Fig. 5). The fluorescence in one cell in the outer supporting cell (DC and HC) area in the cochlear sensory epithelium was bleached by laser zapping (Fig. 5A). After bleaching, fluorescence in the bleached cell gradually recovered as fluorescent dye diffused back from neighboring cells through GJs (Fig. 5A&B). The speed of the FRAP is inversely proportional to GJ permeability. Fig. 5B&C shows that after application of 0.5 mM ACh, the recovery time constant of FRAP was increased to 69.3±5.69 s (n=48, Fig. 5C). In comparison with the time constant (45.8±4.68 s, n=70) in the control group without Ach treatment, ACh significantly increased the recovery time constant of FRAP more than 50% (P<0.001, one-way ANOVA with a Bonferroni correction). That is, ACh diminished GJ permeability between cochlear SCs. Moreover, the effect of ACh on GJ permeability appeared slowly development on the order of minutes (Fig. 5D). The time constant of the effect of ACh on GJ-permeability was around 11.0 min. Glutamate is a major excitatory neurotransmitter in the cochlea and is also thought to be the putative neurotransmitter of the synapses between OHCs and type II ANs (Weisz et al., 2009; 2021). Fig. 5B&C show that application of glutamate (0.2 mM) had no significant effect on GJ permeability between SCs. The time constant of recovery was 42.8±4.65 s (n=32) and had no significant difference from that in the control group (P=0.33, one-way ANOVA).

**Fig. 5.**
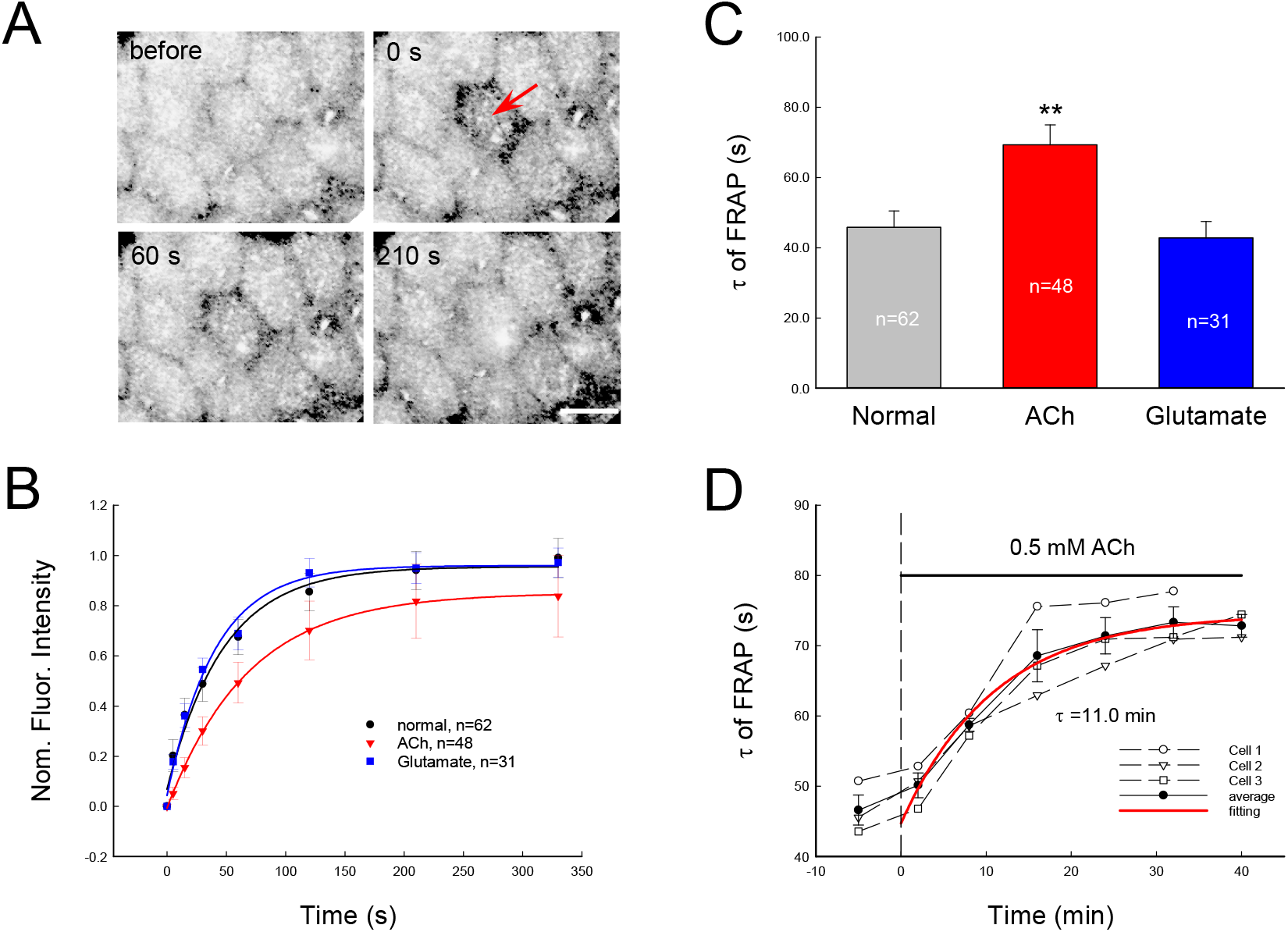
The effect of neurotransmitters of ACh and glutamate on GJ permeability between outer supporting cells (DCs and HCs) measured by FRAP in guinea pigs. **A:** Fluorescent images of cochlear sensory epithelium in the HC area and fluorescence recovery after photobleaching by laser zapping (indicated by a red arrow). Scale bar: 10 µm. **B:** Fluorescence recovery of outer supporting cells at ACh (0.5 mM) or glutamate (0.2 mM) treatment. The data points were averaged from different cells measured at 10 -20 minutes after treatment of ACh or glutamate. Solid lines represent exponential fitting to data. **C:** The recovery time constant (τ) of FRAP. ACh but not glutamate significantly increased the recovery time constant of FRAP, i.e., reduced GJ permeability. **: P<0.001, one-way ANOVA with a Bonferroni correction. **D:** Dynamic changes of GJ permeability in cochlear supporting cells for application of ACh. The time constants of FRAP were measured pre- and post-application of 0.5 mM ACh. The black circles and solid line represent the average value from three measurements. The red line represents exponential fitting. The time constant of the fitting is 11.0 min.

### Effects of ACh and uncoupling of GJs between SCs on OHC electromotility

OHCs have electromotility (Brownell et al., 1985; Zheng et al., 2000), which plays a critical role in the mammalian hearing (Liberman et al., 2002). Fig. 6A-C show the direct effect of ACh on OHC electromotility. After application of 0.1 mM ACh, OHC electromotility associated nonlinear capacitance (NLC) was shifted to the left negative voltage direction. The peak voltage (V_pk_) of NLC was shifted from -22.8±2.50 mV in control level to -30.7±2.92 mV (n=16, P=2.68E-07, pared t test, two-tail) (Fig. 6C). The shift is reversible and repeatable. After stop of application of ACh, V_pk_ returned to pre-application level (Fig. 6B). However, the peak capacitance (C_pk_) had no significant changes. C_pk_ in the control and application of 0.1 mM ACh was 48.2±1.50 pF and 48.4±1.72 pF (n=16), respectively (Fig. 6-S1A). There was no significant difference between them (P=0.71, two-tail, paired t test).

**Fig. 6.**
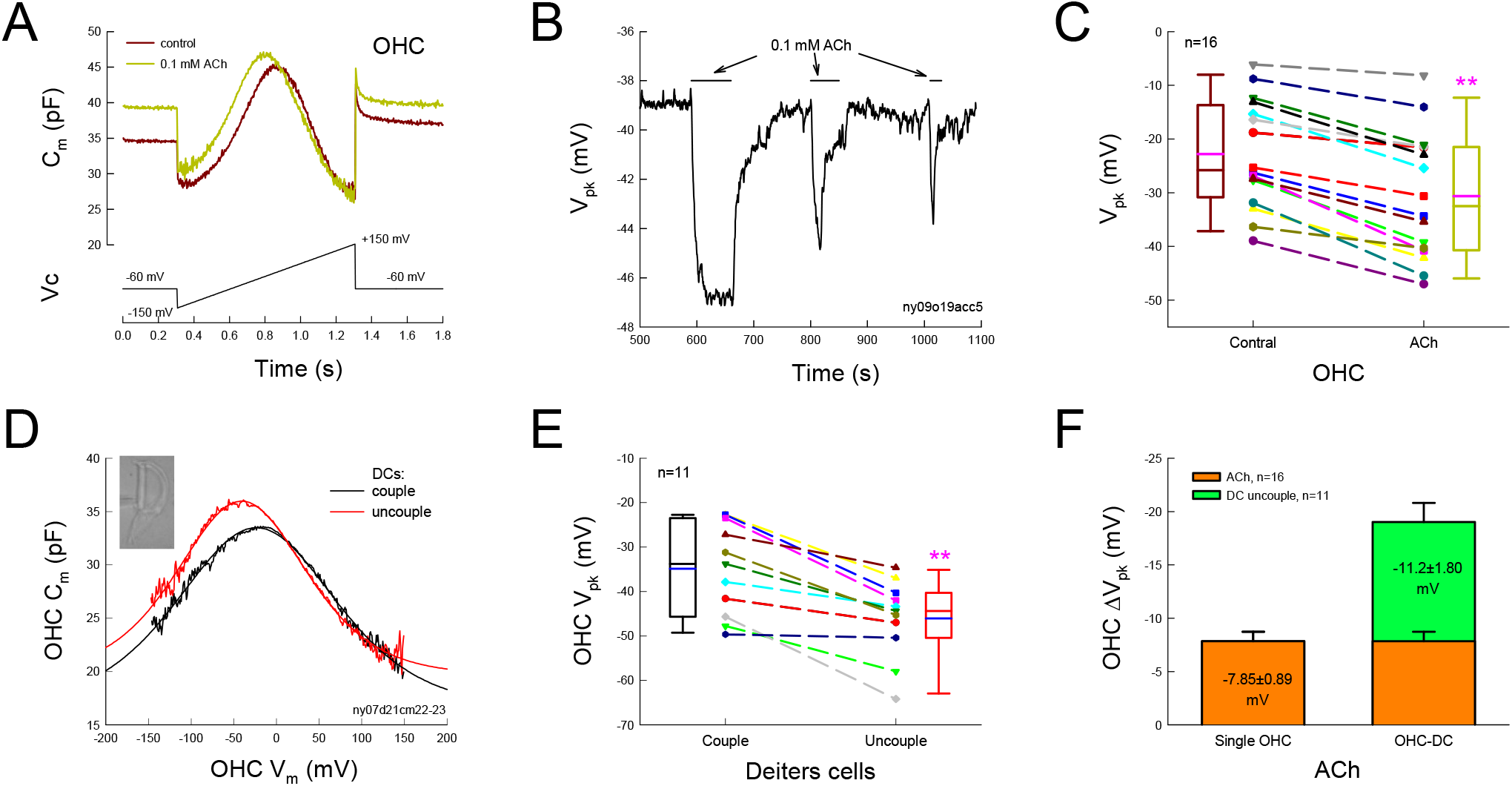
Effects of ACh and uncoupling GJs between DCs on OHC electromotility in guinea pigs. **A-B:** ACh-induced changes in OHC electromotility associated nonlinear capacitance (NLC). After application of 0.1 mM ACh, NLC was shifted to left negative voltage side. Panel B is the continuously tracking the voltage at the peak of NLC (V_pk_). Application of ACh caused V_pk_ shifted to negative voltage. The shift is reversible and repeatable. **C:** ACh-induced V_pk_ changes. Dashed lines connected two same symbols represent the change of V_pk_ in the same OHC before and after application of 0.1 mM ACh. Pink lines in the boxes represent the mean levels. There is significant change in V_pk_ for application of 0.1 mM ACh (**: P=2.68E-7, paired t test, two-tail). **D:** Influence of uncoupling GJs between DCs on OHC electromotility. Uncoupling GJs between DCs shifted OHC NLC to left negative voltage direction. Smooth lines represent NLC fitted by Boltzmann function. The parameters of fitting are Q_max_=4.20 & 3.29 pC, z=0.42 & 0.51, V_pk_=-23.3 & -40.8 mV, and C_lin_=16.6 & 19.7 pF for coupling and uncoupling of GJs between DCs, respectively. **E:** Uncoupling of GJs between DCs shifts OHC NLC to left side. Dashed lines connected two same symbols represent the change of V_pk_ in the same OHC before and after application of 0.1 mM ACh. Blue lines in the boxes represent the mean levels. There is significant change in OHC V_pk_ for uncoupling GJs between DCs (**: P=0.0001, paired t test, two-tail). **F:** Shift of V_pk_ in single OHC and OHC-DC pairs at the application of ACh and uncoupling of GJs between DCs. In OHC-DC pairs, both direct effect of ACh on OHC electromotility and indirect effect of ACh-induced uncoupling between DCs on OHC electromotility are superimposed to shift OHC electromotility to left side.

Our previous study demonstrated that electronic stimulation and GJ changes in DCs could influence OHC electromotility even there is no GJ between DCs and OHCs (Yu and Zhao, 2009). Since ACh could evoke inward currents in DCs and uncouple GJs between DCs (Figs. 2-4), we further investigated the effect of uncoupling of GJs between DCs on OHC electromotility. Fig. 6D&E show that uncoupling of GJs between the DCs in OHC-DC pairs could shift OHC NLC to the left negative voltage direction as the direct effect of ACh on OHC electromotility. V_pk_ of NLC was shifted from -34.9±3.09 mV to -46.1±2.63 mV (n=11) after uncoupled GJs between DCs by using patch pipette breaking one DC in coupled 2DCs (Fig. 6E). The change was significant (P=0.0001, two-tail paired t test). Also, the C_pk_ of OHCs was changed from 44.0±1.65 pF to 45.4±1.76 pF (n=11) after DCs uncoupled (Fig. 6-S1B). However, the change was not significant (P=0.06, two-tail paired t test). Since both ACh and uncoupling of GJs between DCs produced the same left-shift in OHC NLC, Fig. 6F shows this imposed effect. Application of 0.1 mM ACh shifted NLC V_pk_ by -7.85±0.89 mV (n=16) (Fig. 6C) and the uncoupling of GJs between DCs could shift OHC NLC about -11.2±1.80 mV (n=11) (Fig. 6E). Thus, both effects could produce ∼ -20 mV shift of V_pk_ in OHC NLC in OHC-DC pairs (Fig. 6F).

Moreover, since ACh could evoke large inward currents in DCs (Figs. 2-3), which can influence OHC electromotility as well (Yu and Zhao, 2009), we further did double patch clamp recording to simulate the effect of this current change in DCs on OHC electromotility. Fig. 6-S2 shows that OHC and DC in a pair of OHC and DCs were simultaneously recorded by double patch clamp. When the holding current in the DC was changed from -4 to 4 nA, OHC NLC appeared a left-shift (Fig. 6-S2B), as the same as the effect of ACh and uncoupling of GJs between DCs on OHC NLC (Fig. 6).

### Impairment of the regulation of active cochlear amplification and the protection of hearing from noise by deficiency of GJs between DCs

To assess the function of this SC GJ-mediated efferent control *in vivo*, we selectively deleted Cx26 expression in DCs by crossing with Prox1-Cre line (Zhu et al., 2013; Zong et al., 2017). As reported in previous reports (Zhu et al., 2013; Zong et al., 2017), Cx26 expression in the DCs was selectively deleted and there was no apparent hair cell degeneration in this Cx26-Prox1 conditional knockout (cKO) mouse line (Fig. 7A). However, active cochlear amplification measured as distortion product otoacoustic emission (DPOAE) was reduced (Fig. 7B&C). The gain of amplification, which was measured by amplitude of 2f_1_-f_2_ re f_1_ amplitude in DPOAE, was also reduced and the gain regulation assessed by the I/O function was impaired (Fig. 7D). Thresholds of auditory brainstem response (ABR) in Cx26 cKO mice also showed moderate hearing loss, especially at the high frequency range (Fig. 7-S1).

**Fig. 7.**
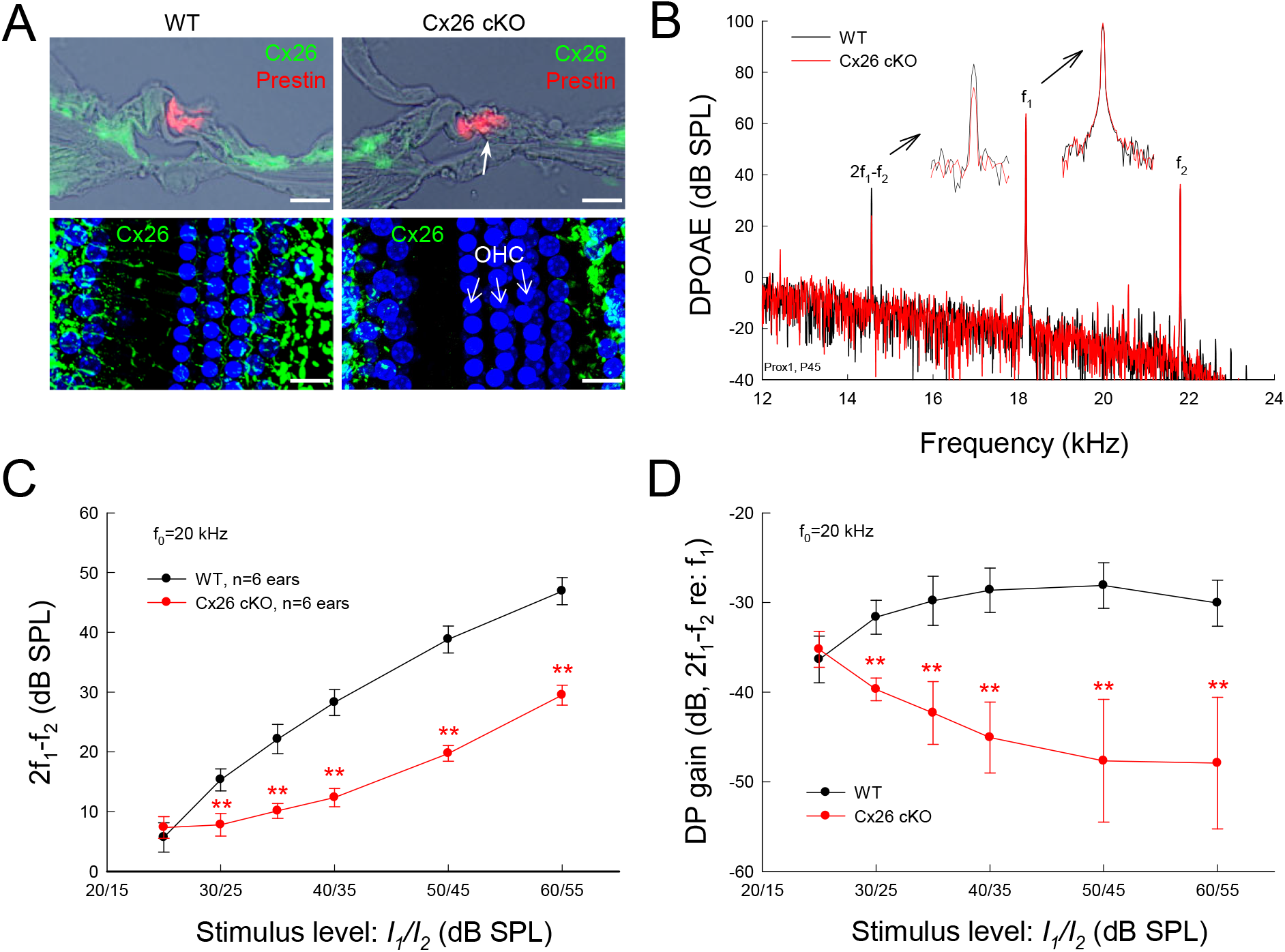
Deficiency of the GJ-mediated efferent pathway by targeted-deletion of Cx26 expression in DCs declines active cochlear amplification and regulation. Wild-type (WT) littermates served as control. Mice were postnatal day 45 old. **A:** Targeted-deletion of Cx26 in DCs. A white arrow indicates lack of Cx26 labeling in the DC area in Cx26 cKO mice in immunofluorescent staining for Cx26 (green). OHCs were visualized by prestin staining (red). Bottom images show whole-mounting view of targeted-deletion of Cx26 in DCs by immunofluorescent labeling for Cx26. Scale bars: 50 µm. **B:** Spectrum of acoustic emission recorded from Cx26 cKO mice and WT mice. Insets: Large scale plotting of 2f_1_-f_2_ and f_1_ peaks. The peak of DPOAE (2f_1_-f_2_) in Cx26 cKO mice was reduced but f_1_ and f_2_ peaks remained the same as those in WT mice. f_0_=20 kHz, I_1_/I_2_=60/55 dB SPL. **C:** Reduction of DPOAE in Cx26 cKO mice in I/O plot. **D:** I-O function of DP gain (2f_1_-f_2_ re: f_1_) in Cx26 cKO mice and WT mice. DP gain in WT mice was almost flat, whereas the DP gain in Cx26 cKO mice decreased as sound intensity increased. **: P < 0.01, two-tail t test.

We further tested the protective function from noise exposure in this mouse line. Fig. 8 shows that after exposure to ∼96 dB wide noise for 2-hr, the WT mice had quickly a recovery in 3 days and completely recovered at post-exposure day 28 measured by DPOAE and ABR recordings (Fig. 8C&D). However, DPOAE in Cx26-Prox1 cKO mice had no recovery after noise exposure (Fig. 8A), and was continuously reduced and reached the maximum reduction at post-exposure day 3 (a blue arrow indicated in Fig. 8A). ABR thresholds in Cx26 cKO mice also had significant increases after noise exposure and did not return to the control levels in comparison with control Cx26 cKO mice without noise exposure at the same age (Fig. 8B). It is worthy to be noted that the control Cx26-Prox1 cKO mice without noise exposure had a progressive hearing loss and DPOAE reduction with age increased (Fig. 8A&B), consistent with our previous report that this Cx26-Prox1 cKO mouse line has progressive hearing loss (Zong et al., 2017).

**Fig. 8.**
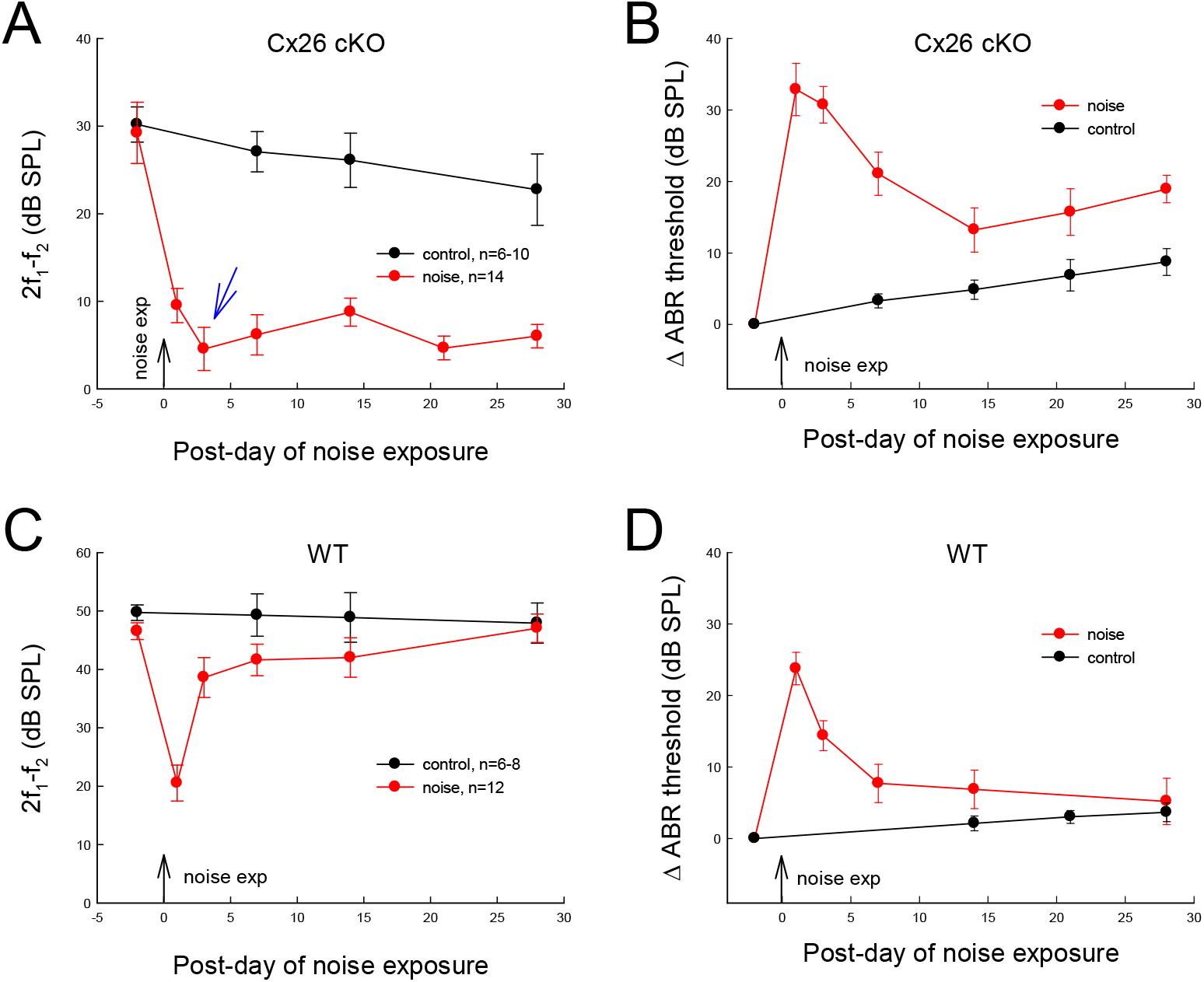
Deficiency of GJ-mediated control pathway increases susceptibility to noise in mice. Mice were exposed to 96 dB SPL while-noise for 2-hr, one time. Black vertical arrows indicate the noise-exposure day, which was defined as post-exposure day 0. Control mice were not exposed to noise. DPOAE (2f_1_-f_2_) was measured at f_0_=20 kHz, I_1_/I_2_=60/55 dB SPL. ABR thresholds were measured by 16 kHz tone-bursts and normalized to the pre-noise exposure level. **A-B:** Noise-induced DPOAE and ABR threshold changes in Cx26 cKO mice. A blue arrow in panel A indicates that DPOAE was reduced to the minimum level at post-exposure day 3 in Cx26 cKO mice. DPOAE and ABR thresholds in Cx26 cKO mice were not completely recovered after noise exposure. **C-D:** DPOAE and ABR threshold changes in WT mice for noise exposure. DPOAE and ABR thresholds were completely recovered at post-exposure day 28.

## Discussion

In this experiment, we found that MOC efferent nerves have functional innervations with SCs in the cochlea (Figs. 1-3). Application of MOC neurotransmitter ACh could evoke inward currents in the SCs and reduced GJs between them (Figs. 2-5). This ACh-induced uncoupling effect on GJs between the SCs had long-lasting influence (Figs. 4&5) and enhanced the direct inhibition of ACh on OHC electromotility (Fig. 6). Deficiency of this SC GJ-mediated control by targeted-deletion of Cx26 between DCs compromised the regulation of active cochlear amplification and the protection of hearing from noise trauma (Figs. 7&8). Taken together, these data demonstrate that the MOC efferent system has functional innervations with the cochlear SCs to regulate GJs between them. This control could enhance the direct modulation of ACh on OHC electromotility and the regulation of hearing sensitivity. These data also suggest that this SC GJ-mediated efferent control has a critical role in the protection of hearing from noise trauma.

Previous studies reported that there are nerve innervations in the cochlear SCs and they could form chemical synapses with outer supporting cells (DCs and HCs) (Wright and Preston, 1976; Stopp and Comis, 1979; Liberman et al., 1990; Nadol and Burgess, 1994; Burgess et al., 1997; Bruce et al., 2000). However, the source, function, and significance of such neural innervations in the cochlear SCs remain unclear or are under debating (Fechner et al., 1998, 2001). In this study, we found that the MOC fibers had branches projecting to the cochlear SCs (Figs. 1 and 1-S1). The SCs also had AChR expression (Figs. 1D&E), which was associated with MOC fibers (Fig. 1-S1). Moreover, presynaptic VAChT labeling was also visible in SCs and associated with MOC nerves (Fig. 1A&B). Finally, application of principle MOC neurotransmitter ACh could evoke current responses in the cochlear SCs (Figs. 2-3). These results also consistent with a previous report that ACh could evoke intracellular Ca^2+^ rising in DCs (Matsunobu et al., 2001). Taken together, these data suggest that MOC fibers have functional innervations in the cochlear SCs.

It has been reported that the cochlear SCs also have innervations of the branches from the type II AN fibers under OHCs (Fechner et al., 1998, 2001; Simmons and Liberman, 1988; Thiers et al., 2002, 2008). However, we found that VAChT and AChR expressions in the SCs were associated with MOC fibers but not associated with eGFP-targeted type II ANs in SCs (Figs. 1A&B and 1-S1). Moreover, the MOC nerves and type II AN fibers showed distinct locations to cross the cochlear tunnel to innervate SCs (Fig. 1A&B). Also, type-II ANs are not cholinergic fibers. Thus, type-II AN fibers are unlikely to be responsible for our observed cholinergic nerve fiber innervations and ACh-evoked responses in SCs.

Application of ACh could evoke inward current in the cochlear SCs (Figs. 2-3). The Hill’s coefficient in the DC was 1.64 (Fig. 2E), which is comparable with ACh-evoked responses in OHCs in previous studies, e.g., 1.6 in Shigemoto and Ohmori (1991), 1.7 in McNiven et al. (1996), and 1.6 in Dallos et al. (1997). The EC_50_ in single DC was 72.6 μM (Fig. 2E), which is higher than that (10-20 μM) recorded in OHCs. However, considering that DCs are electrically coupled by GJs, the electronic response in the DC group could be amplified by 2-3 times by GJ coupling between them (Fig. 3). Thus, the EC_50_ in the group DCs could be reduced by 2-3 times, close to the EC_50_ recorded in OHCs.

The fact that ACh could evoke inward current in the SCs indicates that AChRs in the SCs are nicotinic. It has been reported that nicotinic α9 and α10 subunits are predominant isoforms in the cochlea and are permeable to Ca^2+^ (Elgoyhen et al., 1994, 2001; Vetter et al., 1999, 2007; Weisstaub et al., 2002; Gomez-Casati et al., 2005; Plazas et al., 2005; Lustig, 2006; Lipovsek et al., 2012). We found that SCs had positive labeling to AChRα9 (Fig. 1D&E), consistent with previous reports. A previous study also reported that ACh could elevate intracellular Ca^2+^ concentration in DCs, which could be eliminated by α9 AChR antagonists (Matsunobu et al., 2001). Thus, the SCs may have α9 nAChR expression. However, the detailed information about composition and formation of AChRs in the SCs needs to be further investigated in future.

In this study, we further found that ACh could close GJs between the cochlear SCs (Figs. 4&5). GJ channels, including the inner ear GJ channels, are sensitive to intracellular Ca^2+^; elevation of intracellular Ca^2+^ can close GJ channels (Spray et al., 1982; Sato and Santos-Sacchi, 1994). We previously reported that ATP can activate purinergic P2X receptors leading to Ca^2+^ influx in the cochlear SCs to close GJs between them (Zhu and Zhao, 2012). ACh closing GJs between SCs may take the same mechanism. As discussed above, nicotinic AChRs in the cochlea are formed by α9 and α10 subunits, which are highly Ca^2+^ permeable (Weisstaub et al., 2002; Gomez-Casati et al., 2005; Lipovsek et al., 2012). We found that the SCs had positive labeling to AChRα9 (Fig. 1D&E). Moreover, a previous study demonstrated that the ACh could elevate intracellular Ca^2+^ concentration in DCs (Matsunobu et al., 2001). Thus, ACh could activate ACh receptors in the SCs and influx Ca^2+^ to elevate intracellular Ca^2+^, which may lead to close GJ channels between them.

Furthermore, we found that uncoupling of GJs between DCs could shift OHC electromotility to the left hyperpolarization direction and enhanced the direct effect of ACh on OHC electromotility (Fig. 6). There is no electronic and GJ coupling between DCs and OHCs (Zhao and Santos-Sacchi, 1999; Zhao, 2000). However, *in situ*, phalangeal processes of DCs attach to the apex of an OHC at the reticular lamina and their basal ends extend to the basal of another OHC (inset in Figs. 6D and 6-S2A). Thus, the mechanical changes in DCs can alter OHC loading or tension, which can modulate OHC electromotility due to OHC electromotility is membrane tension- or loading-dependent (Iwasa, 1993; Kakahata and Santos-Sacchi, 1995; Ashmore 2008). On the other hand, the DC has microfilaments along its phalangeal process. It has been reported that Ca^2+^ influx and electronic stimulation can induce the head of the DC phalangeal process a small movement and increase its stiffness (Dulon et al., 1994; Bobbin et al., 2002), which could consequently alter OHC loading or tension and modulate OHC electromotility. Indeed, our previous study demonstrated that the membrane current changes in DCs and uncoupling of GJs between them could modulate OHC electromotility; the effect could be eliminated by destruction of cytoskeleton in DCs (Yu and Zhao, 2009). In this study, we found that ACh could evoke inward current in DCs (Figs. 2-3) and uncouple GJs between them (Figs. 4-5). These changes in DCs may take the same mechanism, i.e., by altering OHC loading or tension to modulate OHC electromotility.

In the study, we found ACh shifted OHC NLC to the left hyperpolarization direction (Fig. 6A-C) but did not reduce NLC (Fig. 6-S1). This seems contrary to the inhibitory effect of the MOC system on the cochlear responses *in vivo*. Indeed, it was reported that ACh increased motile response measured in the isolated OHCs (Dallos et al., 1997). At present, the answer for this puzzle or the detailed mechanism for the MOC efferent system to inhibit active cochlear mechanics *in vivo* is still unknown. Our finding that MOC fibers have innervations in SCs (Fig. 1); ACh could evoke inward current in DCs (Figs. 2-3) and uncoupled GJs between them (Figs. 4-5). ACh also could evoke intracellular Ca^2+^ elevation in DCs (Matsunobu et al., 2001). As discussed above, these ACh-evoked responses could cause the DC phalangeal process contraction and increase its stiffness (Dulon et al., 1994; Bobbin et al., 2002). Due to OHC electromotility is loading or tension-dependent (Iwasa, 1993; Kakahata and Santos-Sacchi, 1995; Ashmore 2008), these contractions could consequently alter OHC-loading to shift OHC electromotility (Fig. 6D&E and also see Yu and Zhao, 2009). In particular, the SCs are extensively coupled by GJs. These ACh-evoked SC contractions may also increase the stiffness of whole organ of Corti. Taken together, these ACh-evoked responses and effects may eventually reduce active cochlear amplification *in vivo*. Indeed, it has been reported that whole organ of Corti including DCs and HCs *in vivo* was contracted to reduce sensitivity under acoustic overstimulation (Fridberger et al., 1998; Flock et al., 1999). Moreover, this contraction could last long time (>30 min) after the stimulation was terminated, suggesting that SCs may take an active part in the protection (Fridberger et al., 1998; Flock et al., 1999). These data also provide further evidence that SCs have the MOC efferent system control and may have an important role in the efferent inhibition of active cochlear amplification *in vivo*.

These data may also have an implication that this SC-mediated efferent control may have a role in the slow MOC effect. The MOC efferent system has a slow effect on the order of minutes in addition of the fast effect (Sridhar et al., 1995; Guinan, 2006). At the present, the mechanism underlying this slow MOC effect still remains unclear. In the experiment, we found that different from the fast dynamic of the direct effect of ACh on OHC electromotility (Fig. 6B), the effect of ACh on GJs between SCs was relatively slow and had long-lasting influence (Figs. 4&5). This is consistent with observation that contraction of SCs lasted longer time even the stimulation was terminated (Fridberger et al., 1998; Flock et al., 1999). In addition, it has been reported that the slow MOC effect plays a critical role in the protection of hearing from noise (Guinan, 2006). In the experiment, we found that impairment of this SC GJ-mediated pathway compromised the regulation of active cochlear amplification and increased susceptibility to noise (Figs. 7&8). These data demonstrated that this ‘indirect’ efferent control via SCs plays an important role in the protection of hearing from noise trauma and also suggested that this SC GJ-mediated efferent control may have a role in the slow cochlear efferent effect.

## Materials and Methods

### Animal selection and preparation

In this study, adult mice (1-6 months old) and guinea pigs (200-450 g) with either gender were used. Most of electrophysiological recording were performed in guinea pigs, since the whole cochlear sensory epithelium (including the basal turn) could be easily isolated and could obtain large numbers of the cochlear SCs and OHCs, in particular, OHC-DC pairs. Most of histological and *in vivo* examinations were performed in mice, since most of transgenic animal models are mice. We also did electrophysiological recording in mice to verify the ACh responses in the cochlear SCs.

Adult Hartley guinea pigs were purchased from (Charles River Laboratories, USA) and used in experiments. For mouse experiments, adult CBA/CaJ mice (Stock No: 000654, The Jackson Lab, USA) were used. Peripherin-eGFP transgenic mice (McLenachan et al., 2008), which were gifted from Dr. Ebenezer Yamoah at University of Nevada, were also used in morphological experiments. For targeted deletion of Cx26 in outer supporting cells, *Cx26^loxP/loxP^* transgenic mice (EM00245, European Mouse Mutant Archive) were crossed with mice of the *Prox1-CreER^T2^* Cre line (Stock No. 022075, Jackson Laboratory, USA). As we previously reported (Zhu et al., 2013; Zong et al., 2017), Tamoxifen (T5648, Sigma-Aldrich, St. Louis, MO) was administrated to all litters at postnatal day 0 (P0) by intraperitoneal injection (0.5 mg/10g x 3 days). WT littermates were used as control. All experimental procedures were conducted in accordance with the policies of the University of Kentucky Animal Care & Use Committee. Either gender of animals was used in the experiments.

### Immunofluorescent staining with confocal microscopy

Immunofluorescent staining was performed as described in our previous reports (Zhao and Yu, 2006; Liu and Zhao, 2008). The mouse cochlea was freshly isolated and the temporal bone was removed. The otic capsule was opened. The round window membrane was broken by a needle and a small hole was also made at the apical tip of the cochlea. Then, the isolated cochlea was emerged into 4% paraformaldehyde for fixation for 30 min. After washout with PBS, the cochlea was decalcified by 10% EDTA for 5-8 hr and the cochlear sensory epithelium was isolated after removing the bone and stria vascularis. After washout with PBS, the isolated epithelia were incubated in a blocking solution (10% goat serum and 1% BSA in PBS) with 0.5% Triton X-100 and then incubated with polyclonal chicken anti-neurofilament (1:500, Cat# AB5539, Millipre Corp, CA), monoclonal mouse anti-Tuj1 (1:500, Cat# MMS-435P, BioLegend, CA), monoclonal chicken anti-GFP (1:500, Cat# ab13970, Abcam, MA), monoclonal mouse anti-AChRα9 (1:100, Cat# sc-293282, Santa Cruz Biotech Inc, CA), monoclonal rabbit anti-VAChT (1:500, Cat# ab279710, Abcam, MA), monoclonal mouse anti-Sox2 (1:200, Cat# sc-365823, Santa Cruz Biotech Inc, CA), polyclonal goat anti-prestin (1:50, Cat# sc-22694, Santa Cruz Biotech Inc, CA), or monoclonal mouse anti-Cx26 (1: 400, Cat# 33-5800, Invitrogen), at 4°C overnight. After being washed with PBS for 3 times, the epithelia were incubated with corresponding Alexa Fluor conjugated second antibodies (1:500, Molecular Probes) at room temperature (23 °C) for 1 hr.

After mounting on the glass slide, the stained epithelia were observed under a Nikon A1R confocal microscope system with Nikon 60x or 100x Plan Apro oil objective. Serial sections were scanned along the z-axis from the bottom to apical surface of the epithelium with a 0.25 μm step. NIS Elements AR Analysis software (Nikon) was used for constructing 3D image from z-stack section.

### Cochlear sensory epithelium and cell isolation for electrophysiological recording

As we previously reported (Zhao and Yu, 2006; Yu and Zhao, 2008, 2009; Zhu and Zhao, 2010, 2012), guinea pigs or mice were decapitated and the temporal bone was removed. The otic capsule was dissected in the normal extracellular solution (NES, 142 NaCl, 5.37 KCl, 1.47 MgCl_2_, 2 CaCl_2_, 10 HEPES in mM, 300 mOsm and pH 7.2). After removal of the bone and stria vascularis, the sensory epithelium (organ of Corti) was exposed and picked away along the whole cochlea with a sharpened needle. For electrophysiological recording, the isolated sensory epithelia were further dissociated by trypsin (0.5 mg/ml) for 2-3 minutes with shaking. Then, the dissociated cells and small cell groups were transferred to the recording chamber. The cochlear OHCs and SCs could be unequivocally identified under microscope by their own morphological shape (Zhu and Zhao, 2010, 2012).

### Patch-clamp recording

The isolated cells were continuously perfused with the NES (0.5 mL/min). The selected OHC or SC was recorded under the whole-cell configuration using an Axopatch 200B patch clamp amplifier (Molecular Devices, CA, USA) (Yu and Zhao, 2009; Zhu and Zhao, 2010). Patch pipettes were filled with an intracellular solution (140 mM KCl, 5 mM EGTA, 2 mM MgCl_2_, and 10 mM HEPES, pH 7.2, 300 mOsm) with initial resistance of 2.5-3.5 MΩ in bath solution. Data were collected by jClamp software (SciSoft, New Haven, CT, USA). The signal was filtered by a 4-pole low-pass Bessel filter with a cut-off frequency of 2 kHz and digitized utilizing a Digidata 1322A (Molecular Devices, CA, USA). All recordings were performed at room temperature (23°C) and finished within 4-6 hrs after isolation to ensure cells having good conditions. The recording was stopped when the seal resistance became worse <500 MΩ in the SC recording.

*C_in_* was continually recorded online at 1-3 Hz from the transient charge induced by small (-10 mV) test pulses with duration of 18X the time constant at the holding potential. The transient charge was calculated from the integration of capacitance current with time (Zhao and Santos-Sacchi, 1998). Membrane potential (*V_m_*) was corrected for pipette series resistance (*R_s_*).

The OHC electromotility associated NLC was measured with a two-sinusoidal wave voltage stimulus in jClamp (Yu and Zhao, 2008, 2009). This voltage stimulus was composed of a ramp command (-150 mV to +150 mV) summed with two sinusoidal commands (f_1_=390.6 Hz, f_2_=781.3 Hz, 25 mV peak to peak). The signal was filtered by a 4-pole low-pass Bessel filter with a cut-off frequency of 10 kHz. The capacitance was calculated by admittance analysis of the current response. In some cases, the peak of NLC and the voltage corresponding to the peak capacitance (*V_pk_*) were continuously recorded by a tracking technique (sampling rate: 4/s, tracking-step: 0.25 mV) (Zhao et al., 2005; Yu and Zhao, 2008, 2009).

Data analysis was performed with jClamp and MATLAB (Zhao and Santos-Sacchi, 1999; Yu and Zhao, 2008, 2009). The voltage-dependent NLC was fitted to the first derivative of a two-state Boltzmann function:

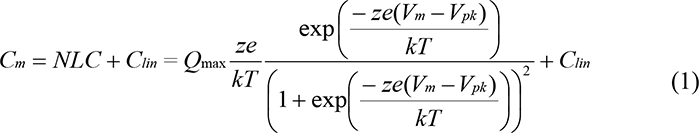

where *Q_max_* is the maximum charge transferred, *V_pk_* is the potential that corresponds to the peak of NLC and also has an equal charge distribution, *z* is the number of elementary charge (*e*), *k* is Boltzmann’s constant, *T* is the absolute temperature and *C_lin_* is the cell membrane capacitance. Curve fitting and figure plotting was performed with SigmaPlot software. Membrane potential (*V_m_*) was corrected for pipette series resistance (*R_s_*).

For double patch clamp recording in DC-OHC pairs, a pair of 2DCs connected 1 or 2 OHCs isolated from the guinea pig cochlea was selected. One pipette was patched at the DC and another patch pipette was patched at the basal nuclear pole of the OHC under a whole-cell configuration by using Axopatch 700A (Molecular Devices, CA, USA). The command and data recording in each patch clamp were separately controlled by jClamp (SciSoft, New Haven, CT, USA).

### Fluorescence recovery after photobleaching (FRAP)

The freshly isolated guinea pig cochlear epithelium or cells were incubated in 10 µM Carboxy SNARF-1 AM (C-1271, Molecular Probes) at room temperature for 15-30 min, protected from light. Then, the incubated tissues or cells were continuously perfused with the NES for 30-45 min to remove the residual dye and allow completion of the hydrolysis of AM ester prior to measurement.

FRAP measuring was performed by use of a laser scanning confocal system. Prior to laser bleaching, the fluorescence image of the selected area in the cochlear sensory epithelium was scanned and saved. Then, a laser beam (the laser power was nominally about 50 µW) illuminated a selected cell for 20 seconds to bleach the fluorescence. After bleaching, the fluorescent images were taken at 0, 5, 15, 30, 60, 120, 210 and 330 s to measure the recovery. The intensity of fluorescence of the bleached cell was measured off-line by use of ImageJ software (NIH, Bethesda, MD) and normalized to the fluorescent intensity in the same cell prior to bleaching. The fluorescence recovery data were fitted by:

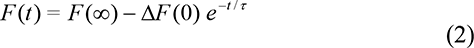

where *F(∞)* is the fluorescence signal of the bleached cell at *t* = ∞, Δ*F(0)* is the initial fluorescence change due to the bleaching, and *τ* is the recovery time constant.

### Noise exposure

Mice were awake in a small cage under loud-speakers in a sound-proof chamber and exposed to white-noise (96 dB SPL) for 2 hr, one time. Sound pressure level and spectrum in the cage were measured prior to placement of the animal (Zhao et al., 2021).

### ABR and DPOAE recording

As we previously reported (Zhu et al., 2013, 2015), the ABR was recorded by use of a Tucker-Davis ABR & DPOAE workstation with ES-1 high frequency speaker (Tucker-Davis Tech. Alachua, FL). Mice were anesthetized by intraperitoneal injection with a mixture of ketamine and xylazine (8.5 ml saline+1 ml Ketamine+0.55 ml Xylazine, 0.1 ml/10 g). Body temperature was maintained at 37–38°C. ABR was measured by clicks and tone bursts (8 – 40 kHz) from 80 to 10 dB SPL in a 5 dB step. The ABR threshold was determined by the lowest level at which an ABR can be recognized. If mice had severe hearing loss, the ABR test from 110 to 70 dB SPL was added.

DPOAE was recorded as described by our previous reports (Zhu et al., 2013, 2015). Two pure tones (f_1_ and f_2_) were simultaneously delivered into the ear through two plastic tubes coupled to two high-frequency speakers (EC-1, Tucker-Davis Tech. Alachua, FL). The test frequencies were presented by a geometric mean of f_1_ and f_2_ [f_0_ = (f_1_ x f_2_)^1/2^] from f_0_=4 to 20 kHz. The ratio of f_2_ versus f_1_ (f_2_/f_1_) was 1.22. The intensity of f_1_ was set at 5 dB SPL higher than that of f_2_. One hundred fifty responses were averaged. A cubic distortion component of 2f_1_-f_2_ in DPOAEs was measured.

### Chemicals and data processing

All chemicals were purchased from Sigma Chemical Company (St. Louis, U.S.A.). Chemicals were delivered by a Y-tube perfusion system (Yu and Zhao, 2008, 2009; Zhu and Zhao, 2010). Patch clamp data analyses were performed by jClamp and MATLAB (Zhao and Santos-Sacchi, 1999; Yu and Zhao, 2008, 2009). FRAP data analyses were performed with MATLAB. Data were expressed as mean ± s.e.m. unless otherwise indicated in text and plotted by SigmaPlot (SPSS Inc. Chicago, IL).

### Statistical analysis and reproducibility

The statistical analyses were performed by SPSS v18.0 (SPSS Inc. Chicago, IL). Parametric and nonparametric data comparisons were performed using one-way ANOVA or Student t tests after assessment of normality and variance. The threshold for significance was P= 0.05. Bonferroni *post hoc* test was used in ANOVA.

The numbers of recording times in each experiment were indicated in figures. For *in vivo* noise exposure experiment, only one or two mice were exposed to noise at each time. Totally, 7 mice were exposed in 4 times of noise exposure.

## Acknowledgments

We thank Dr. Michael Bennett for valuable comments on the earlier version of this manuscript. This work was supported by NIH R01 DC 017025, R01 DC 019687, and R56 DC 016585 to HBZ.

## Author Contributions

HBZ conceived the general framework of this study. YZ, LML, LM, YN, CL, JC, YMZ, and HBZ performed the experiments. HBZ, NY, YZ, and LML analyzed data. HBZ wrote the paper. All authors reviewed the manuscript and provided the input.

---

**Fig. 1-S1.**
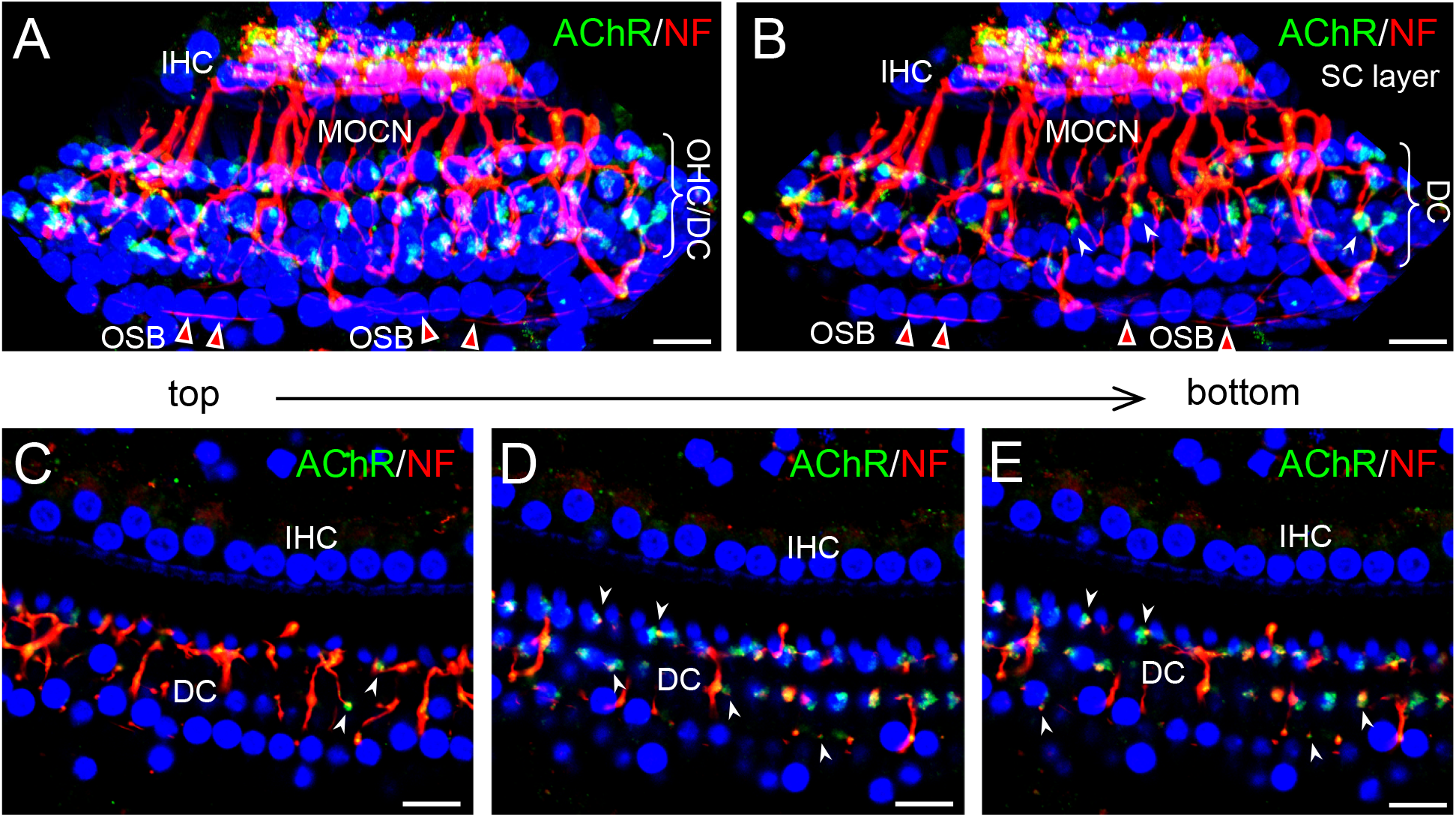
Immunofluorescent staining for AChR (green) and neurofilament (NF, red) in the mouse cochlear sensory epithelium with whole-mount preparation. **A-B:** Top-surface view of 3D images of the cochlear sensory epithelium constructed from z-stack of serial confocal scanning. Panel **B** represents the view at the SC layer after removing OHC layer in the 3D image. White arrowheads indicate synaptic-like structures with AChR and NF labeling. Triangles with red color indicate outer spiral bundle (OSB) of type-II ANs running among SCs. **C-E:** Confocal images were obtained from scanning at different depths in the SC layer. White arrowheads indicate synaptic-like structure with AChR and NF labeling. DC: Deiters cell, IHC: inner hair cell, MOCN: medial olivocochlear nerve. Scale bar: 20 µm

**Fig. 2-S1.**
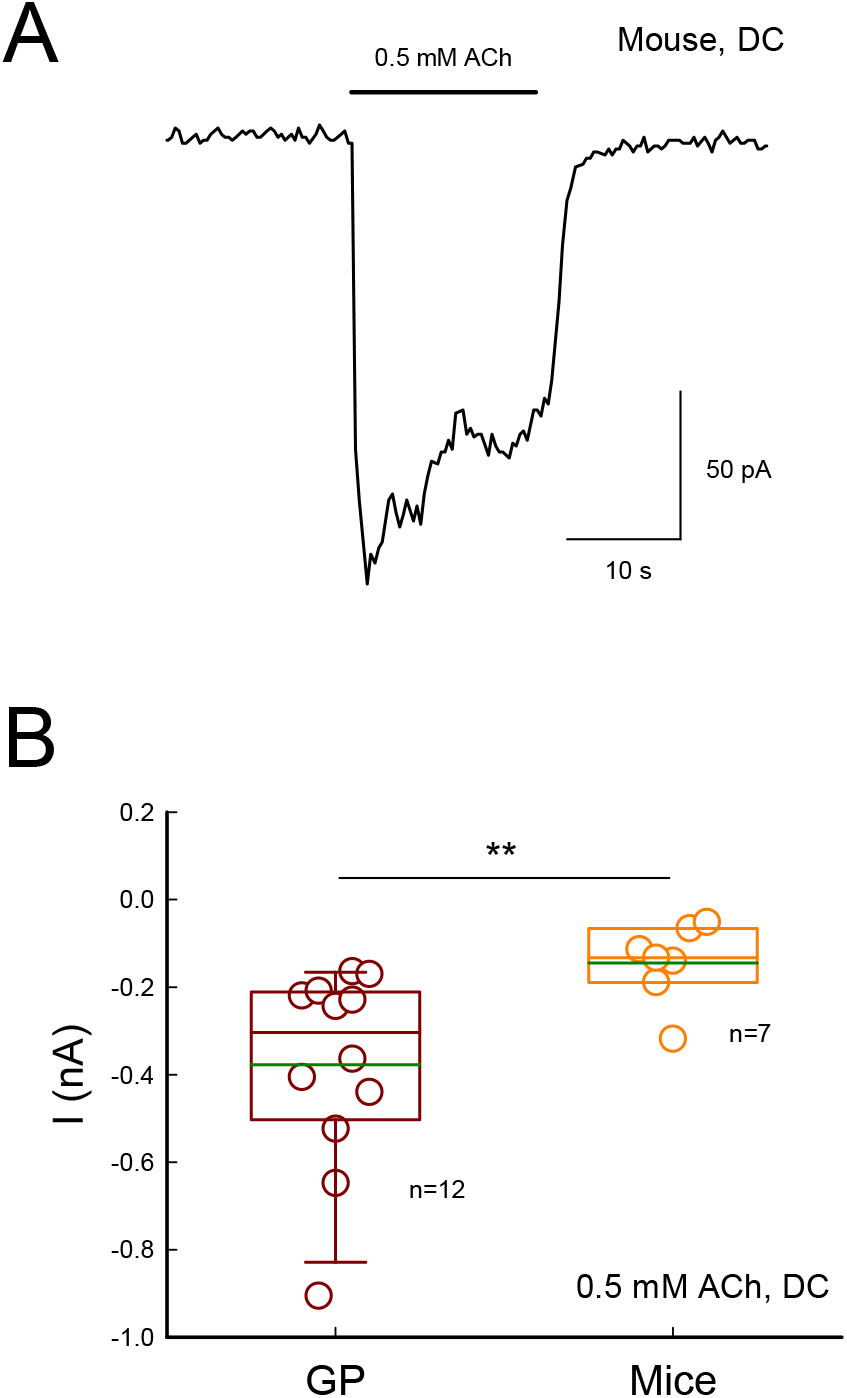
ACh-evoked inward currents in mouse DCs. **A:** ACh-evoked current in single mouse DC. **B:** Comparison of ACh-evoked currents in DCs in guinea pigs and mice. The green lines in boxes represent the mean levels. The average currents evoked by 0.5 mM ACh in guinea pig and mouse DCs are -0.38±0.07 and -0.14±0.03 nA, respectively. The evoked current in mice is significantly smaller than that in guinea pigs (P=0.006, t test, two-tail).

**Fig. 2-S2.**
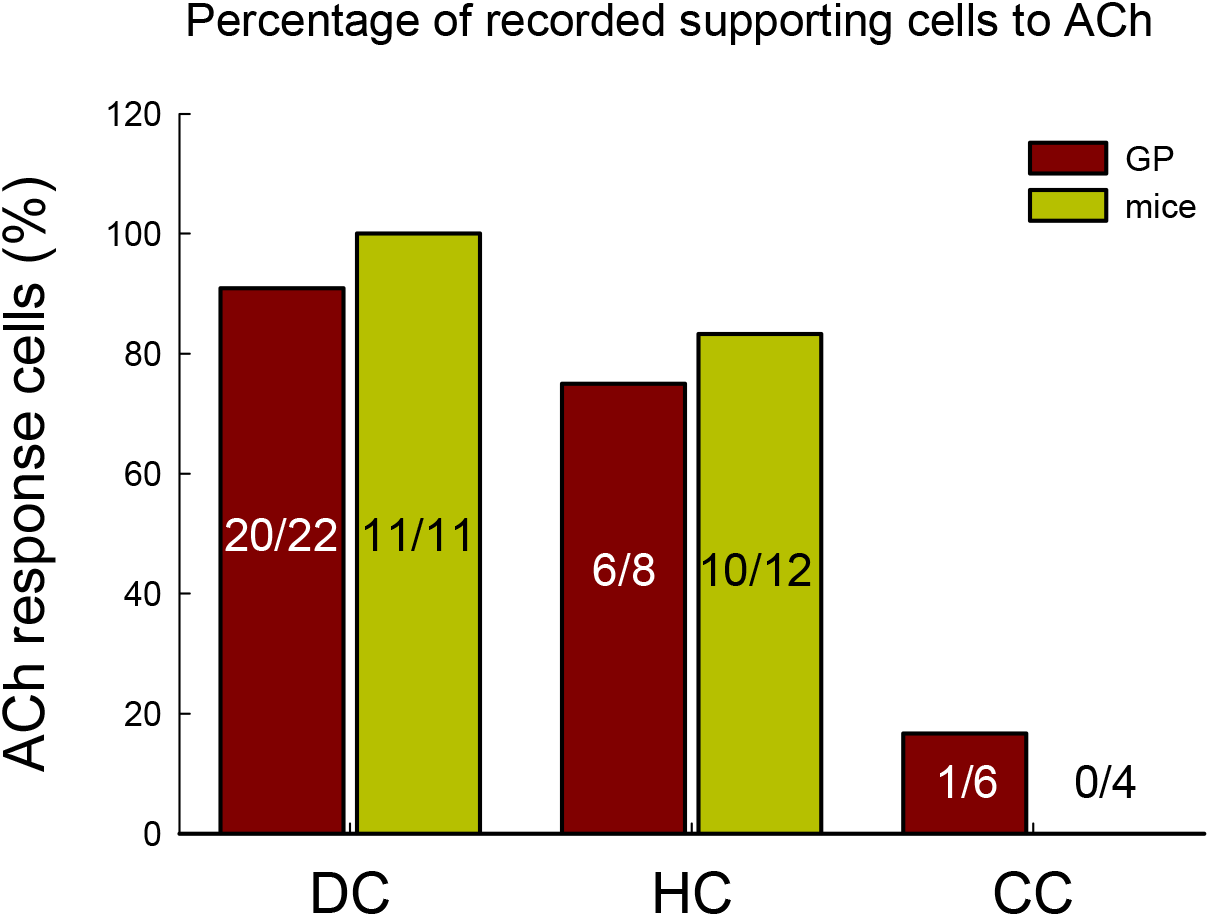
Percentage of ACh-evoked responses in the recorded supporting cells in guinea pigs and mice. Numbers within each bar represent the number of cells with ACh-evoked responses *vs* total of the recorded cells. In both guinea pigs and mice, 91-100% of recorded Deiters cells (DCs) and 75-83% of recorded Hensen cells (HCs) could be evoked responses by ACh. However, only few Claudius cells (CCs, 1/6) in guinea pigs and none of the recorded mouse CCs (0/4) have responses to 0.5 mM ACh.

**Fig. 4-S1.**
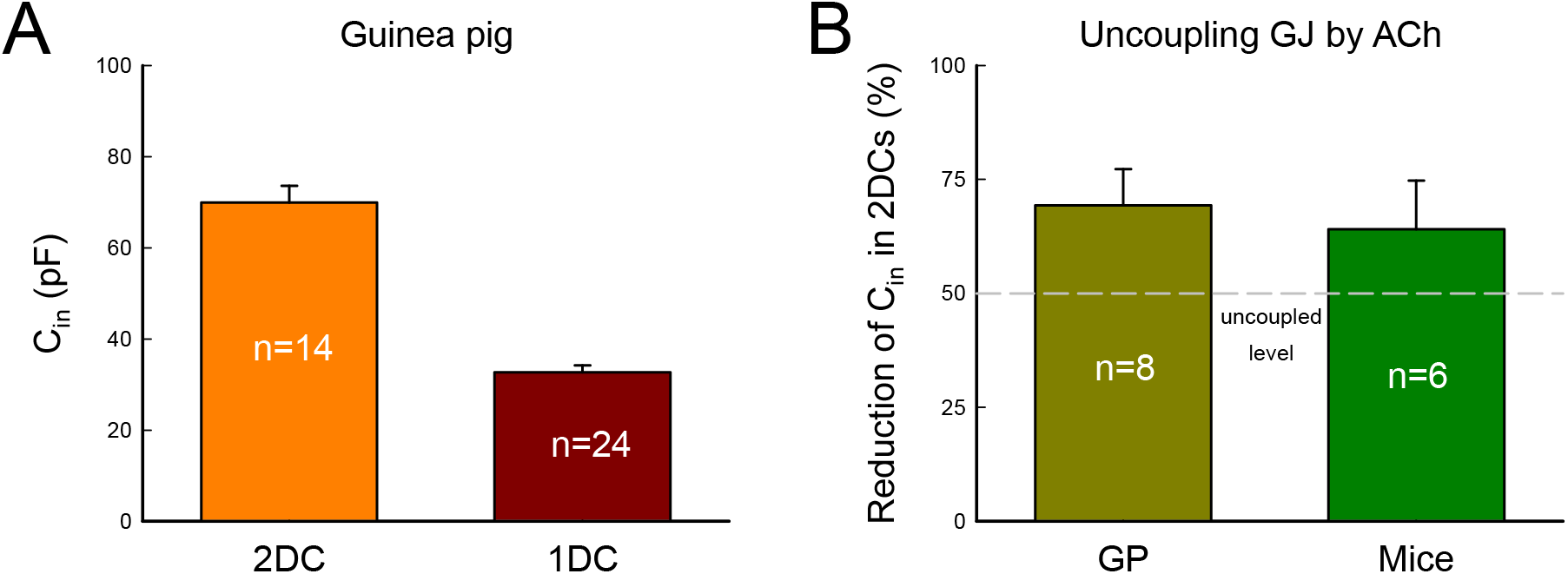
C_in_ in 2DC and 1DC in guinea pigs and reduction of C_in_ in coupled SCs by ACh. **A:** C_in_ measured in two-coupled DCs and single DC in guinea pigs. C_in_ of 2DCs and single DC in guinea pigs is 69.9±3.62 pF and 32.6±1.54 pF, respectively. C_in_ of single DC is about half of C_in_ of 2DCs. **B:** Uncoupling of GJs by ACh. ACh (0.5 mM) reduced C_in_ of guinea pig and mouse 2DCs to 69.3±7.94 and 64.0±10.7%, respectively. The gray dashed line represents the reduction of C_in_ (50%) in 2-coupled cells to the completely uncoupled (one cell) level. There is no difference in the reduction of C_in_ between guinea pigs and mice (P=0.70, t test, two-tail) by ACh.

**Fig. 6-S1.**
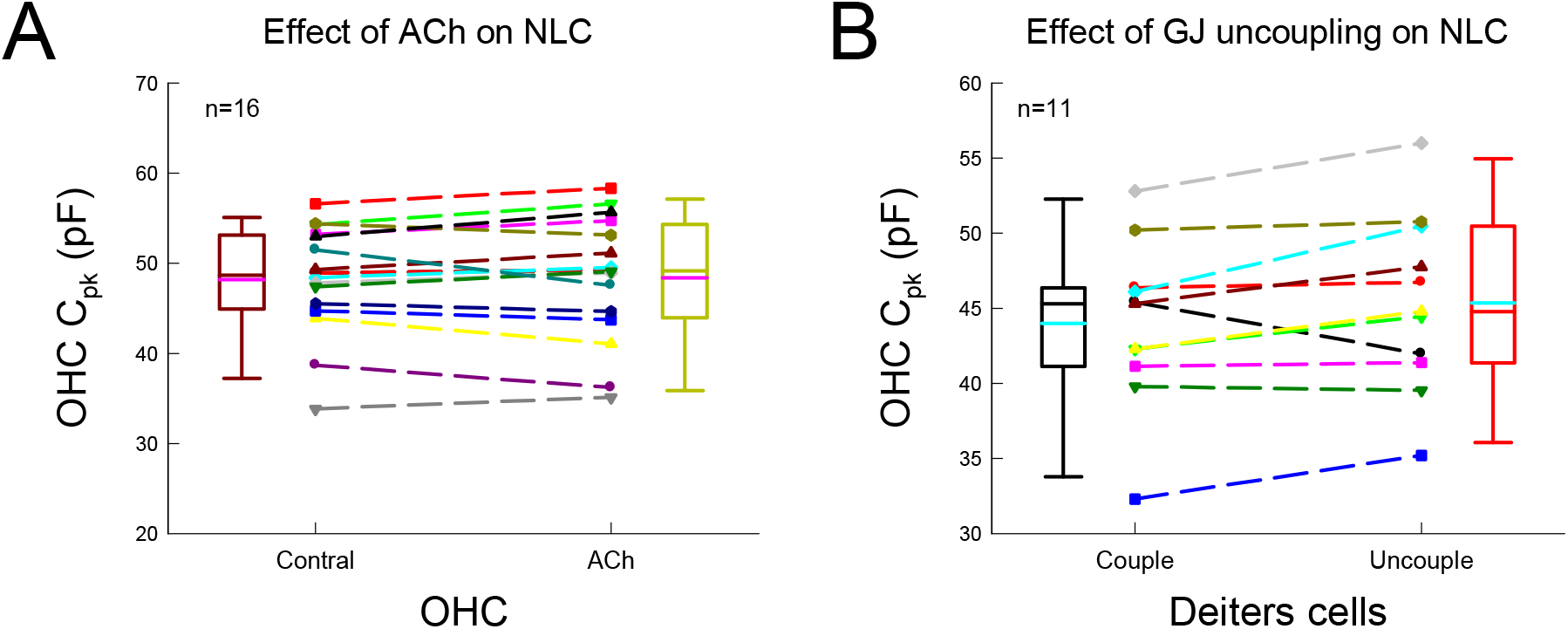
No significant changes in peak capacitance (C_pk_) of OHCs for application of ACh and uncoupling GJs between DCs. **A:** No significant changes in C_pk_ of OHCs for application of 0.1 mM ACh. Dashed lines connected two same symbols represent the change of C_pk_ in the same OHC before and after application of 0.1 mM ACh. Pink lines in the boxes represent the mean levels. There is no significant change in C_pk_ for application of 0.1 mM ACh (P=0.71, paired t test, two-tail). **B:** There is no significant change in OHC C_pk_ in the OHC-DC pair after uncoupled GJs between DCs by mechanical breaking one DC. Dashed lines connected two same symbols represent the C_pk_ change in the same OHC before and after uncoupled GJs between DCs. Cyan lines in the boxes represent the mean levels. There is no significant change in OHC C_pk_ after uncoupled GJs between DCs (P=0.06, paired t test, two-tail).

**Fig. 6-S2.**
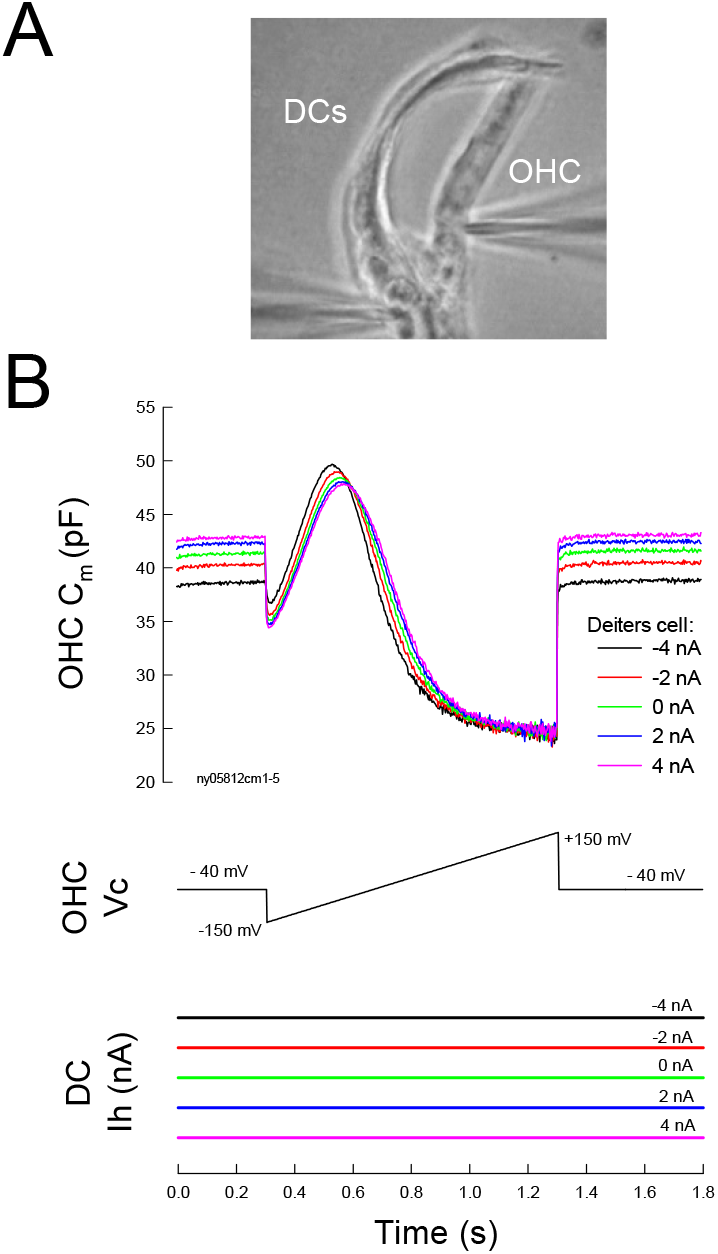
The effect of membrane current changes in DCs on OHC electromotility in a DC-OHC pair. **A:** A captured image of dual-patch clamp recording at a pair of DCs and OHC. One patch pipette was recording at an OHC and another pipette was patched at DCs. **B:** Change of OHC electromotility by alternation of holding currents in DCs. DCs were held at different holding currents and OHC electromotility associated NLC was simultaneously recorded by a voltage ramp with sinusoidal voltage. OHC NLC was left-shifted with injection of negative current in DCs. V_pk_ of NLC was -86.0, -81.0, -77.5, - 74.8, and -72.5 mV for DCs holding at -4, -2, 0, 2, 4 nA, respectively.

**Fig. 7-S1.**
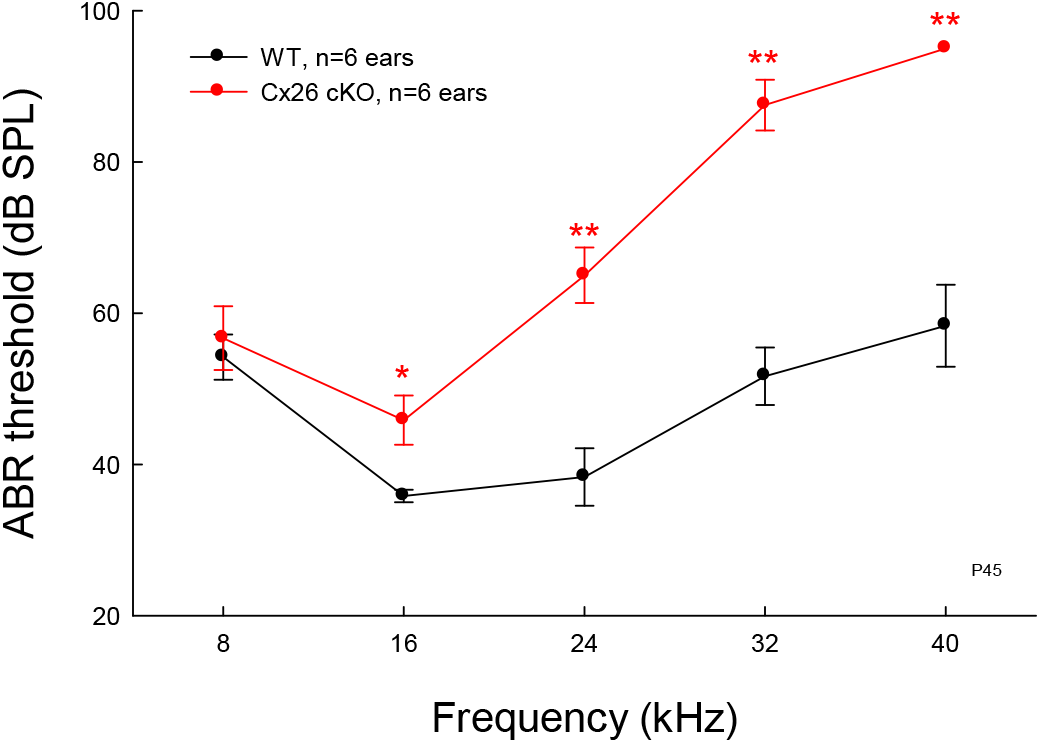
Reduction of hearing sensitivity in Cx26 cKO mice. Mice were 45 days old. WT littermates serve as control. ABR threshold in Cx26 cKO mice was significantly increased. *: P<0.05, **: P<0.01, two-tail t test.

